# Simulation-Based protein engineering of *R. erythropolis* NADH-FMN Oxidoreductase

**DOI:** 10.1101/601252

**Authors:** Ramin Fallahzadeh, Bijan Bambai, Nasrin Kamali, Kasra Esfahani, Abbas Akhavan Sepahi

**Affiliations:** Islamic Azad University Tehran North Branch, Tehran, Iran; National Institute of Genetic Engineering and Biotechnology (NIGEB), Tehran, Iran; Malek Ashtar University of Technology. Tehran, Iran

**Keywords:** *Rhodococcus erythropolis* IGTS8, DszD enzyme, Molecular dynamics simulations, Site-directed mutagenesis, Specific activity

## Abstract

The sulfur contents of fossil fuels has negative impacts on the environment and human health. The bio-catalytic desulfurization strategies and the biological refinement of the fossil fuels are a cost-effective process compared to chemical desulfurization. *Rhodococcus erythropolis* IGTS8 is able to extract the organic sulfur of oil as mineral salts by 4S metabolic pathway (i.e. DszA,B,C and D genes). *dszD* gene codes a NADH:FMN oxidoreductase delivering reducing equivalent to DszA, DszB, and DszC to remove sulfur from heterocyclic molecules. In this study, we sought to improve DszD specificity by side-direct mutagenesis based on prediction of the DszD structure, molecular docking, and molecular dynamic simulation. Accordingly, the cloning, expression and activity assay of best candidates (A79I and A79N) were performed. Our results demonstrated the important role of position 79 in enzyme activity, the A79I and A79N mutants are able to increase the enzyme activity 3.4 and 5.2 fold compared to wild-type.

## Introduction

Biodesulfurization is an attractive alternative process to improve the quality of the refined oil. Removal of sulfur compounds in fossil fuels, especially crude oil, impose an economic and environmental pressure on refinery facilities around the world (Kilbane II 2006). Currently, the method of choice for sulfur reduction at the industrial level is Hydrodesulfurization (HDS). High pressure and temperature along with metal catalysts are used in this process with high cost and some other disadvantages including incapability to decrease all sulfur contents, especially heterocyclic organic compound (Nuhu 2013). In addition, combustion of fossil fuels releases sulfur oxides which accounts for environmental pollution and acid rain (Gupta et al. 2005).

Regarding strict environmental regulations, refinery facilities have been obliged to standardize their products (Akhtar et al. 2016). *Rhodococcus erythropolice* (ATCC 53968) removes up to 900 fold of sulfur from carbon skeleton without any carbon contents reduction. To enhance the bacterial desulfurization rate, in addition to media optimization, one of the key factors is to manipulate the genes involved in biodesulfurization pathway to increase the yield and rate of biodesulfurization process to comparable levels with current chemical processes, but with substantially lower environmental impacts. The preferred method for desulfurization by this microorganism is the 4S metabolic pathway in which DszABCD enzymatic system is used (Davoodi-Dehaghani et al. 2010). DszD is an NADH-dependent FMN oxidoreductase that supplies more FMNH_2_ to both monooxygenases, DszA and DszC (Sucharitakul et al. 2014). DszD is a key enzyme with 192 residues and a molecular weight of ~22 kD. The workflow of this study briefly was divided into four sections, which were:

1. S3-tructure prediction of the DszD enzyme.
2. The binding sites determination of the substrates FMN and NADH on the DszD enzyme.
3. The point mutagenesis in the key residue of the binding site and the molecular assessment by docking and molecular dynamic simulation to select the best mutants for desulfurization application.
4. The cloning, expression and activity assay of the candidate mutants.

## Material & Methods

### Structural prediction and molecular docking

Due to the unavailable crystallographic structure of DszD enzyme, a three dimensional structure predicted for the enzyme through the CPH-models server, was use for structural analysis. This server is a protein homology modeling server (Nielsen et al. 2010). The amino acids sequence of DszD was obtained from NCBI protein sequence database (UniProt: 068503). In this study, homology modeling was applied for structural prediction and recognition of FMN and NADH binding sites in the enzyme. In homologues identification procedure, the sequences with highest identities were selected as templates in the protein data bank (PDB), for structure prediction. These templates revealed the superimposable active sites with DszD by using the Chimera software version 1.10.1. The Chimera software was employed for visualizing the templates and the predicted model (DszD). Furthermore, the 4xj2 protein showed the most superimposable active site among all templates. Overall, 19 conformations mutants obtained using Swiss-PDB Viewer software (Kaplan and Littlejohn 2001) at key residues (i.e. at Thr62, Ser63, Asn77, and Ala79). Then, all mutant and the wild-type optimized for further investigation through the docking tool, Autodock 4.2 (Morris et al. 2009).

### Molecular dynamics simulation of enzyme-substrates complex

After molecular docking, all selected enzyme–substrates complexes (i.e. Enzyme–FMN-NADH) were prepared to be studied through the MD simulations. Simulations were carried out using the GROMACS 5.1.2 package (Abraham et al. 2015) and GROMOS53A6 force field (Oostenbrink et al. 2004). The Automated Topology Builder (ATB) server (Koziara et al. 2014) also provided GROMOS53A6 force field parameters for FMN-NADH molecules. All the systems were first solvated in SPC/E water model (Mark and Nilsson 2001) and neutralized with counter ions in cubic boxes with the length of 7.7 nm. The Periodic Boundary Conditions (PBC) were applied to all axes, short-range electrostatic and van der Waals cut-off set to 1.2 nm in energy minimization of all the systems using the steepest descent algorithm, LINCS algorithm (Hess 2008) also used for holonomic constraints of all bonds in NVT and NPT equilibration of the systems. The particle mesh Ewald (PME) method (Darden et al. 1993) applied in treating long-range electrostatic interactions. The Berendsen thermostat (Berendsen et al. 1984) also applied for temperature coupling at 310K and Parrinello-Rahman algorithm (Parrinello and Rahman 1981) for pressure coupling at 1 bar in the uniform scaling of box vectors (isotropic). Tree independent simulations were performed for 50 ns in this study and further results were analyzed using GROMACS tools as well as MMPBSA method (Kumari et al. 2014) in the calculation of the components of binding energy.

### MMPBSA method in calculation of the free binding energy

The g_mmpbsa package was used to calculate the total free binding energy of the DszD enzyme and its mutants in complex with FMN-NADH. The total binding free energy ΔG_total_ is obtained by calculation of polar interactions binding energy (ΔG_polar_) and non–polar interactions binding energy (ΔG_nonpolar_). The equation 1 demonstrates this calculation.

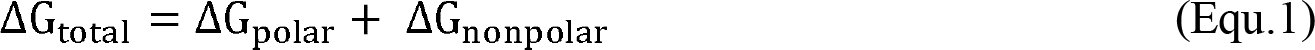

Where each term is obtained by following equations,

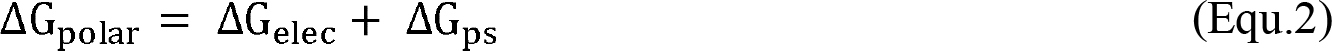

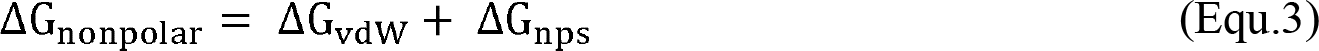

The calculation of energies was performed for the last 30 ns of all MD simulations. The length of the simulation steps was set to 0.2 ns, i.e. 150 snapshots were used totally in the energy calculations (Kumari et al. 2014).

### Studying the stability of secondary structure

Molecular Dynamics Simulation as a potent tool in computational biophysics provides a scientist to gain molecular insights into the dynamical and functional structures as well as the mechanism of action of biomolecules in complex systems. Thus, in order to assess the activity of the enzyme and also its mutants in complex with the substrates in the molecular level we conducted molecular dynamics simulation along molecular docking results. The root mean square deviation (RMSD) analysis performed to investigate the equilibrium state of the wild-type and mutant enzymes in complex with the substrates. The root mean square fluctuation (RMSF) analysis conducted for all the residues in each simulation complex. The dictionary of the secondary structure of protein (DSSP) analysis is also performed to assess the impact of mutations on the entire structure of the native enzyme (Abraham et al. 2015).

### Hydrogen binding analysis

The total number of hydrogen bonds also calculated during the 50 ns of simulation time (Abraham et al. 2015).

### Experimental procedures (benchtop operation)

#### Chemicals, Bacterial strains, plasmids, and Primers

The DszD gene was previously cloned in our lab (Kamali et al. 2010). The cloning vector and host were pBluescript II KS (+) vector (Fermentas) and *Escherichia coli* DH5a cells (Novagen). Furthermore, the expression of wild-type and mutant enzymes were done by using the pET-23a (+) (Novagen) and *E. coli* BL21(DE3) (Promega). DNA polymerase, *Bam* HI and *Eco* RI, T4 Ligase, Ampicillin from Roche Diagnostic (Germany); and DNA ladders, protein markers, T4 DNA ligase and Mouse anti his tag antibody were purchased from Fermentas company. NADH and FMN sodium salts, and the IPTG- X-Gal were purchased from Sigma company. The DNA extraction kit and high pure plasmid purification kit were obtained from Qiagen company. The agarose gel extraction kit and the PCR product purification kit were obtained from Roche company. The LB medium containing 100 μg/ml ampicillin was used in the benchtop operations (at 37 °C).

#### Construction of expression plasmids

In plasmid construction step, the amplicon (599 bp) was cloned using *Bam* HI and *Eco* RI sites of the pET-23a (+) expression vector, named pET-dszD plasmid. Then, the *E. coli* BL21(DE3) cells were transformed with pET-dszD. The dszD mutants were created using the SOEing-PCR method (overlap extension method) (Young et al. 2017). The mutagenic primers and the pET-dszD plasmid were applied as template in SOEing-PCR method. All molecular biology methods were according to the standard protocol (Sambrook et al. 1989).

#### Expression of recombinant genes and analysis

The expression vector was pET-23a that was under control of T7 promoter. Furthermore, the expression was done in *E. coli* BL21(DE3) cells. The original start codon of DszD (TTG) was replaced by ATG of *E. coli* in the beginning of the forward primer in all genes (Table 3). The single colonies were selected from wild-type and mutant plates to examine their expression by induction 1 mM IPTG in LB broth medium for 3 h at 37 °C (OD_600nm_ of 0.6–0.7). The cells harvested by centrifugation and disrupted by ultra-sonication following suspension in Tris-HCl buffer. The lysates centrifuged at 3200 ×g for 5 minutes to remove cell debris and supernatants were used for further analysis. Then, the discontinuous SDS-PAGE was performed under reducing conditions with a %13 polyacrylamide gel (Schägger 2006).

#### Enzyme assay

The assay based on the oxidation of NADH can determine the activity of NADH-FMN oxidoreductase (Kamali et al. 2010). The enzyme activities were determined at 25 °C by quantifying the decrease in absorbance (at 340 nm). The oxidation of NADH to NAD decreased the optical density. The solution of 50 mM Tris/HCl, pH 7.5, 140 μM NADH, 20 μM FMN and varying amounts of cell extract in 1 ml were used as assay mixture. A unit activity is the required amount of flavin reductase to oxidize 1 μM NADH per min (ε_340_ = 6.22 * 10^3^ M^−1^ cm^−1^). The equal concentration of total proteins was used for determination of enzyme activity. The enzymatic activities of samples were compared together.

## Results & Discussion

For enhancing the biodesulfurization capacity, the genes manipulation on the 4S biodesulfurization pathway is necessary. We sought to find key residues in DszD as targets for protein engineering using i*n silico* methods. In this approach, we successfully predict the critical residues involved in the catalytic activity of the DszD enzyme by bioinformatics tools. Before doing the benchtop operation in this study, the bioinformatics workflow briefly was divided into three sections, which were:

1. The crystallographic structure determination of the DszD enzyme.
2. The binding sites determination of the substrates FMN and NADH on the DszD enzyme.
3. The point mutagenesis in the key residue of the binding site and the molecular assessment by docking and dynamic simulation.

### Bioinformatic tools for structural prediction

The crystallographic structure of the DszD enzyme has not yet been determined. Hence, the 3-dimentional structure of this enzyme was determined by the free homology modeling online server CPH in the first step. The tertiary structure of DszD was predicted according to the crystal structures of 4XJ2, representing as the best template molecule. Amino acids sequences of the homologous flavin reductases were aligned with DszD amino acid sequences and revealed that the template have highest primary structure similarity to DszD enzyme. On the other hand, structural data on this enzyme (complex with FMN substrate) were already available at protein data bank (www.rcsb.org). The structural superimposition of the homologue onto the predicted structure of the target revealed that the active center of templates and the target enzyme are similar.

The predicted structure file was named as DszD.pdb. The amino acids sequence of the DszD enzyme (UniProt: 068503) was scanned in the protein data bank for finding the most homologous structures. 22 homologous proteins were found and multiple alignments of these files was performed with the DszD.pdb. Among these aligned sequences, the sequence containing FMN and NADH substrates were selected. Interestingly, a conserved sequence was emerged as the FMN and NADH binding sites in each group of proteins were aligned. By comparing the homologous structures with the DszD structure, the R39, T62, S82, H154, Y179, G182 residues and T62, S63, N77, A79 residues were identified as the FMN and NADH binding sites, respectively.

The superimposition of the active sites of the predicted DszD enzyme with the homologous proteins shows high identities in the software Chimera. Moreover, the closest residues to the FMN substrate were Thr62, Ser63, Asn77, and Ala79 (Fig. 2). Regard to the previous study (Kamali et al. 2010) the substitution of Thr62 by Asn and Ala increased the reaction rate. We conducted this *in silico* study to assess the significance of mutating the critical residues residing in binding site in enzyme activity and structural stability.

Finally, nineteen candidate mutants were created by the SPDB viewer software in these key residues (i.e. at the T62, S63, N77 and A79 residues).

### Model preparation and molecular docking

After energy minimization, both FMN and NADH substrates were docked into wild-type enzyme and these created mutants in the Autodock software. In Figure 1 the wild-type enzyme successfully docked to both NADH and FMN substrates. Table 1 demonstrates the binding energy in the selected mutations. Regarding docking scores, 2 conformations of mutant enzymes presented the lowest binding energies compared to the free energy of wild-type enzyme in binding with FMN-NADH. These 2 mutants include a single substitution of Alanine 79 by Asparagine (A79N mutant) and Isoleucine (A79I mutant). Hydrogen bond lengths between acceptor and donor atoms of the wild-type and mutants and FMN molecule are also indicated in Table 2 which presents the considerable number of hydrogen bonds between the proteins and the substrate.

**Table 1.** Molecular Docking results of the wild-type enzyme and mutants in complex with the substrate FMN

| NO | enzymes | Lowest Binding energy | Mean binding energy |
| --- | --- | --- | --- |
| 1 | wild-type DszD | -8.35 | -8.25 |
| 2 | T62A | -8.65 | -8.58 |
| 3 | T62N | -8.99 | -8.87 |
| 4 | T62S | -8.73 | -8.62 |
| 5 | T62Q | -8.52 | -8.48 |
| 6 | S63A | -8.33 | -8.27 |
| 7 | S63T | -8.54 | -8.48 |
| 8 | N77A | -8.40 | -8.33 |
| 9 | N77C | -8.35 | -8.26 |
| 10 | N77F | -8.40 | -8.33 |
| 11 | N77G | -8.31 | -8.21 |
| 12 | N77S | -8.40 | -8.36 |
| 13 | N77V | -8.46 | -8.39 |
| 14 | A79Q | -8.94 | -8.82 |
| 15 | A79S | -8.43 | -8.37 |
| 16 | A79V | -8.70 | -8.66 |
| 17 | A79T | -8.83 | -8.76 |
| 18 | A79I | -9.07 | -8.99 |
| 19 | A79N | -9.83 | -9.77 |

**Table 2.**
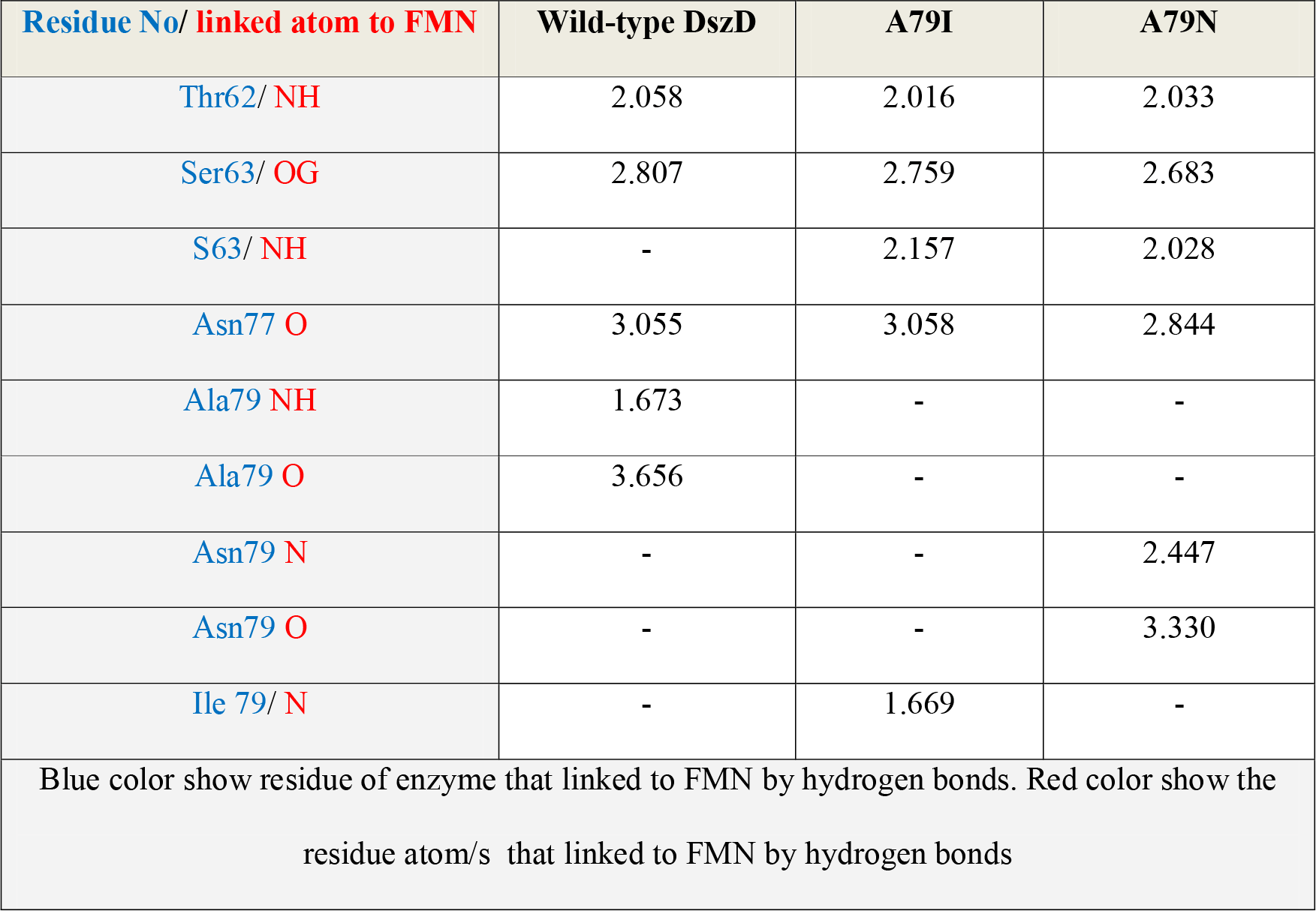
Hydrogen bond lengths between acceptor and donor atoms of the wild-type and mutants and FMN molecule

**Table 3.**
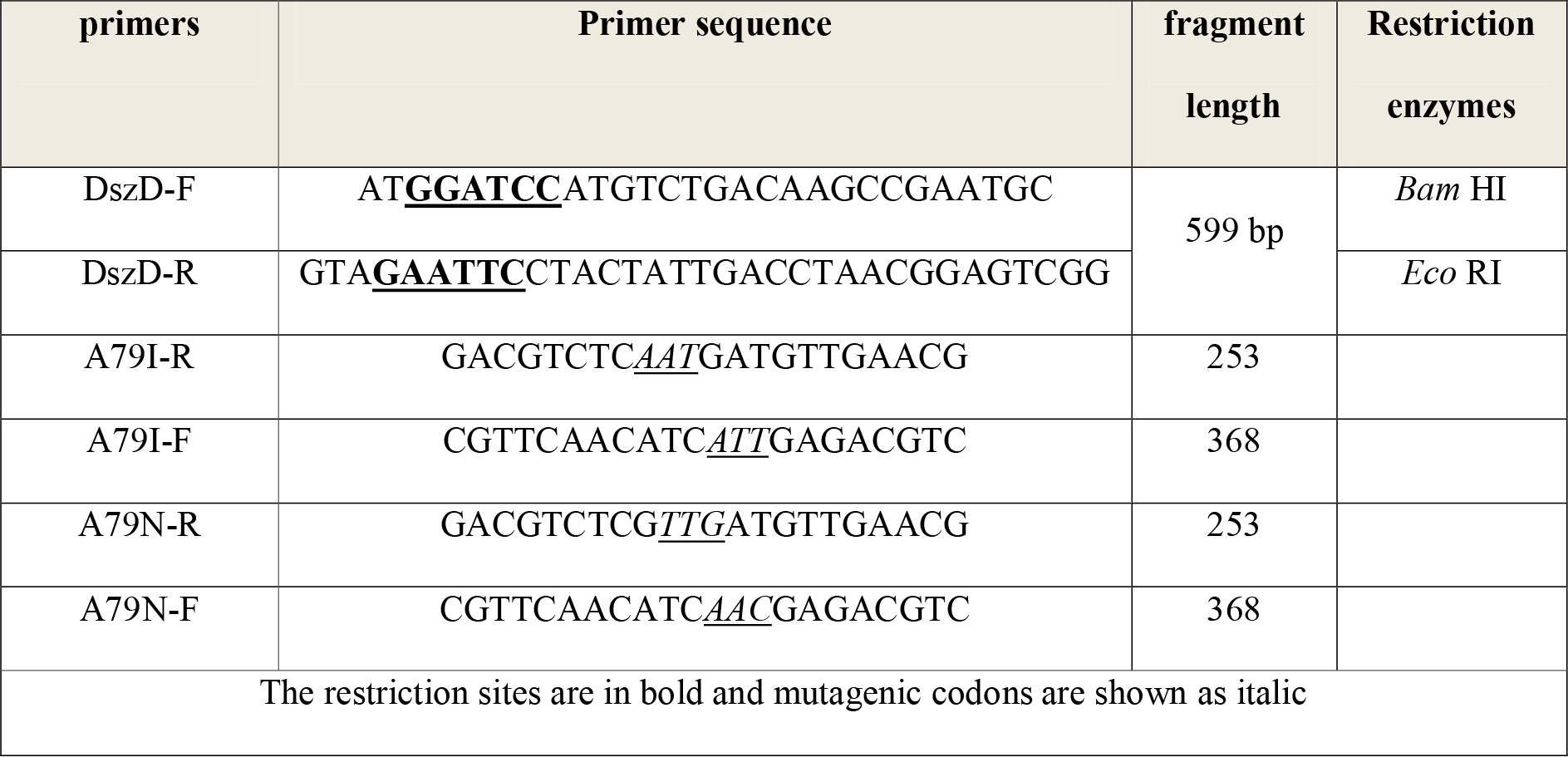
Wild-type and mutagenic primer sequences

**Fig. 1.**
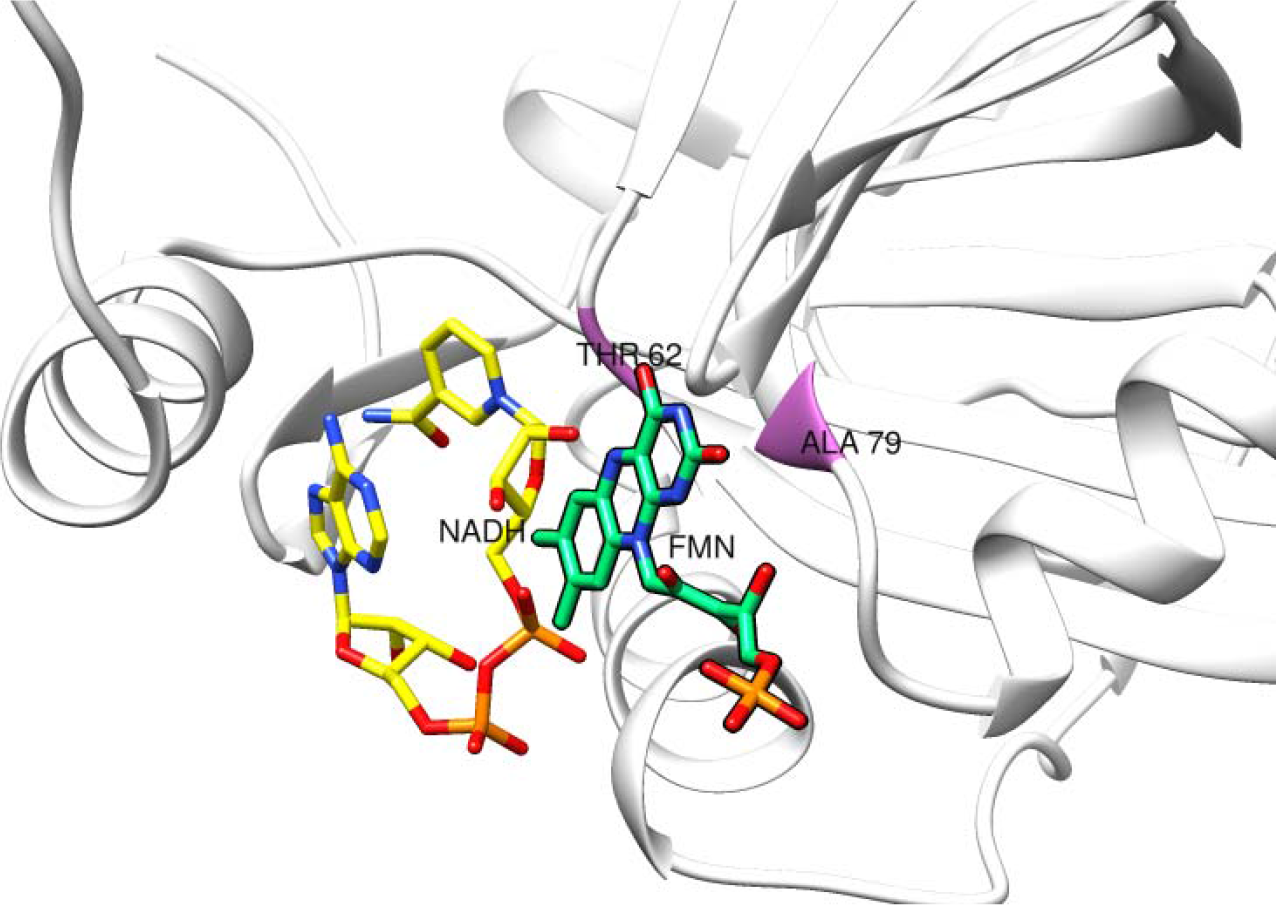
Schematic view of the predicted three-dimensional structure of the wild-type DszD enzyme conformation docked to FMN-NADH substrates

**Fig. 2.**
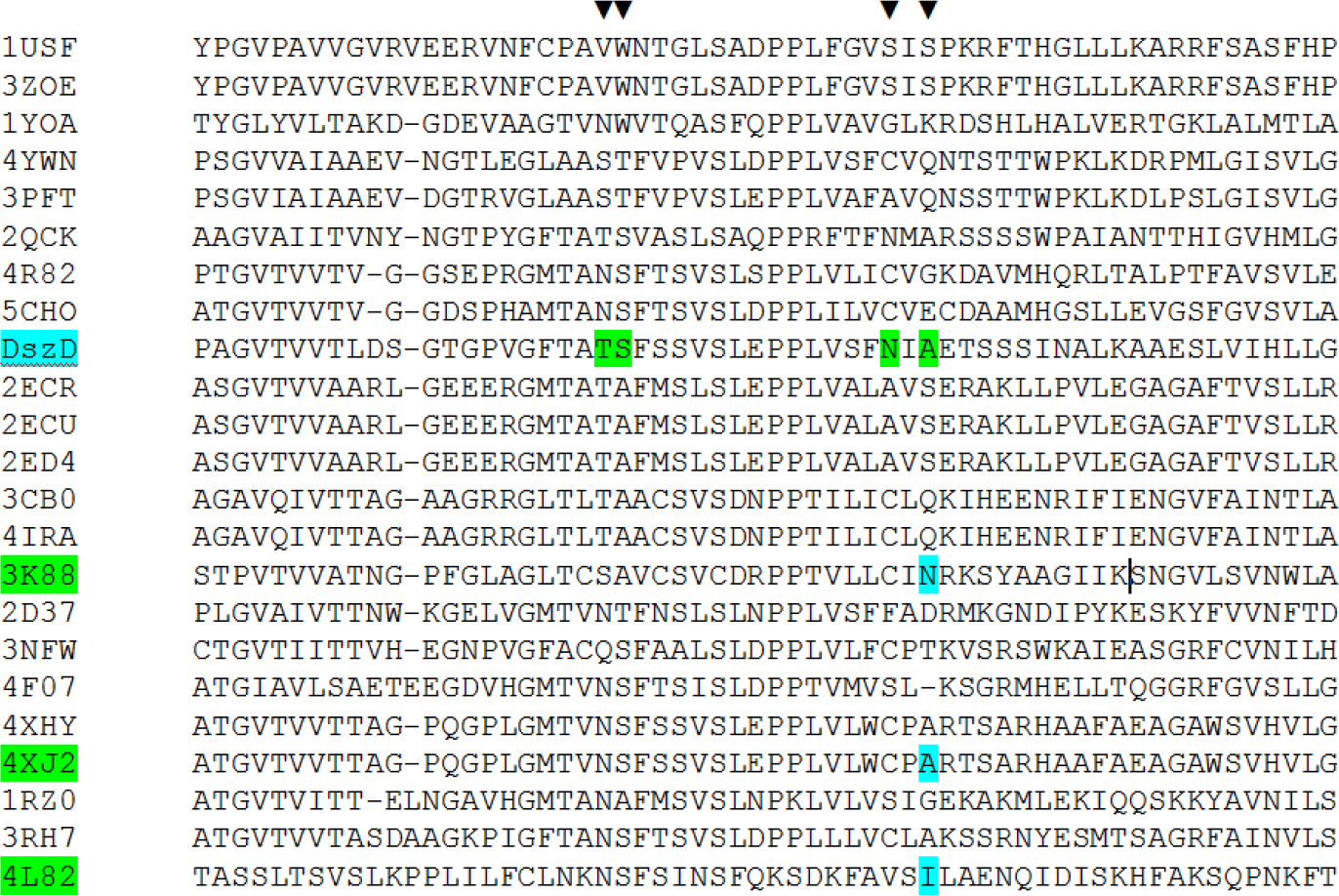
The primary alignment of wild-type DszD enzyme with homologues proteins. In all homologues, FMN bonds to the corresponding residues T62, S63, N77 and A79 of the wild-type (DszD). The Asparagine (N), Alanine (A) and Isoleucine (I) were corresponded with the A79 residue of the wild-type in 3k88, 4xj2 and 4L82 homologues, respectively.

It was very interesting that the similar substitutions (i.e. A79 residue exchange to N or I residue) were presented on the homologous proteins 3K88 and 4L82, respectively. Figure 2 demonstrates key residues (T62, S63, N77 and A79) for binding of substrate FMN. Accordingly, the Ala79 residue of the target enzyme was more confirmed to substituted with asparagine (A79N mutant) or Isoleucine (A79I mutant), corresponding to 3K88-Asn106, and 4L82-Ile82, respectively. The alanine 79 position in DszD enzyme is the amino acid candidate for mutation.

The deduced modeling analysis verified significant conformational space, topology and steric orientation similarities between predicted structure for target DszD-substrates complex and each of template homologs 4XJ2, 3K88 and 4L82. Also the observations of computer analysis supporting that Ala79 is located at 1.67 Å radius isoalloxazine ring-O_2_ atom of substrate FMN. Other important attributes based on primary sequence alignments defined 4XJ2-Ala37 residue, 4L82-Ile82 and 3K88-Asn79 correspondence to DszD-Ala79.

In the 3K88 and 4L82 homologue proteins, the O_2_ atom of the isoalloxazine ring is important in functioning as a recipient of the anion from NAD(P)H.

### Molecular dynamics simulation of enzyme-substrates complex

These data were further confirmed with MD simulation at an atomic level. To study the impact of mutations on DszD structure and DszD-FMN interaction, we performed All-Atom molecular dynamics (MD) simulation. In this study, 50 ns of MD simulation performed on each wild-type and mutant DszD forms in complex with FMN and NADH.

### The free binding energy

Regard to the calculation results of the total free binding energy in each simulation, mutant A79N showed the minimum energy in the enzyme-substrate complex that was represented in Figure 3. The Ala79 substitution by Asparagine and Isoleucine gave favorable binding energies, however, the substitution of Isoleucine in Ala79 position could not justify a favorable mutation due to the large positive number of binding free energy. As demonstrated in hydrogen binding analysis section, a large number of hydrogen bonds in the wild-type enzyme may not necessarily lead to a large negative binding free energy due to the presence of large positive value of polar solvation in equation 2 which affects the total binding free energy of the wild-type enzyme adversely. As indicated in Figure 3 the mutant enzyme A79N offers the most favorable binding energy thus the contribution of this substation to the total binding free energy is also calculated and demonstrated in Figure 4 Consistent with experimental results in this study, substitutions at position 79 by Asparagine and Isoleucine, respectively, favored the binding of mutant enzyme A79N and A79I to the FMN substrate.

**Fig. 3.**
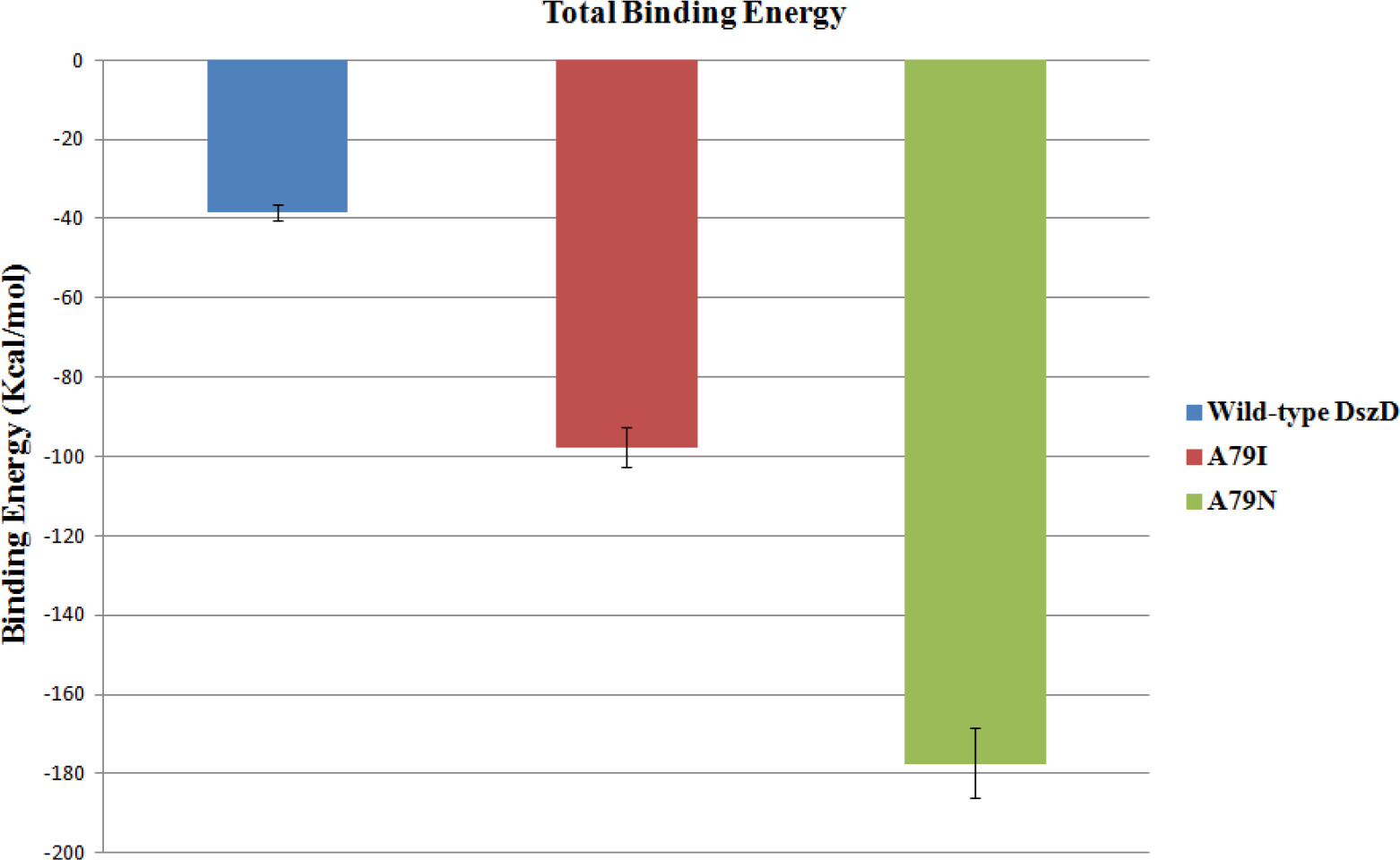
Free Binding energy calculation for the enzyme-substrate complexes.

**Fig. 4.**
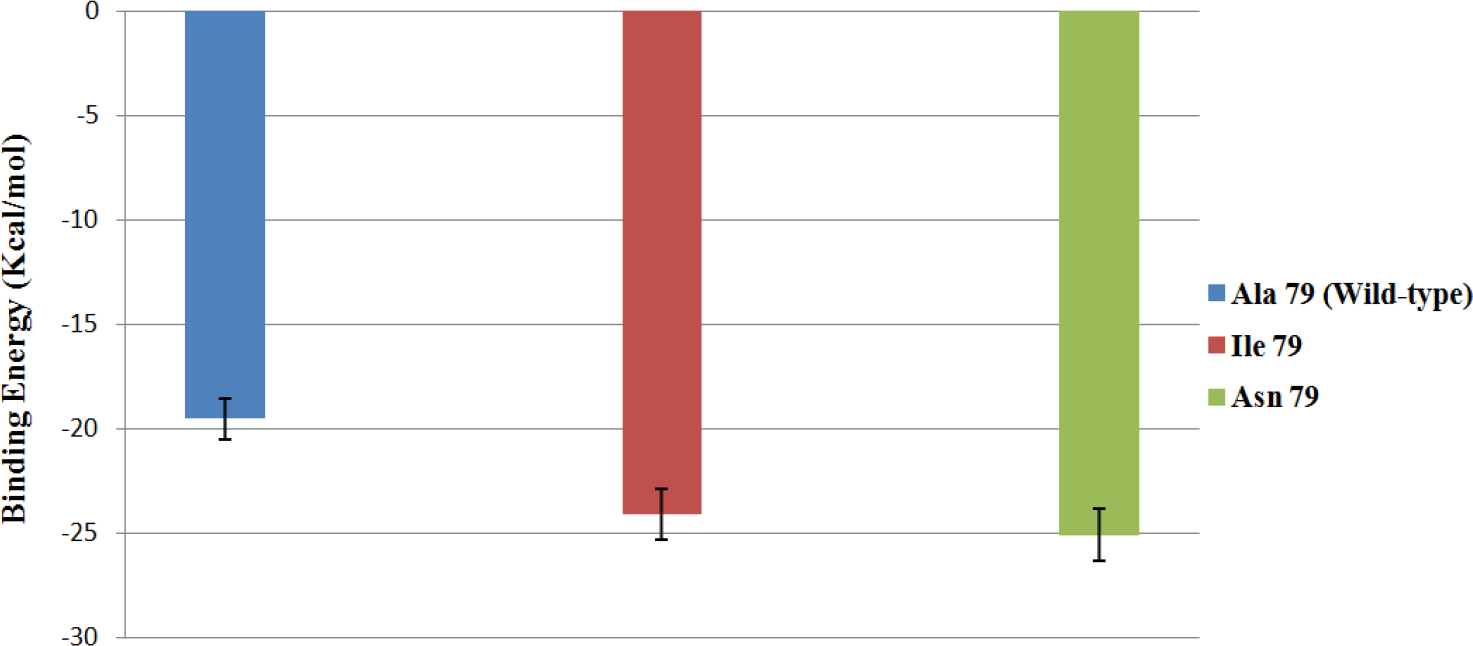
The contributions of critical residues in the transformation of the wild-type enzyme to the mutant enzyme A79I and A79N, in the total binding free energy

### Studying the stability of secondary structure

Figure 5 show the RMSD analysis of the enzyme in wild-type and mutant structures. In all simulations, the enzymes reach a plateau in 20 ns of simulation time and exhibit less conformational changes in last 10 ns of simulation time. The average of RMSD in mutant structures and the native structure show in Figure 6. This was obvious that the stability of mutant did not decrease compared to native structure.

**Fig. 5.**
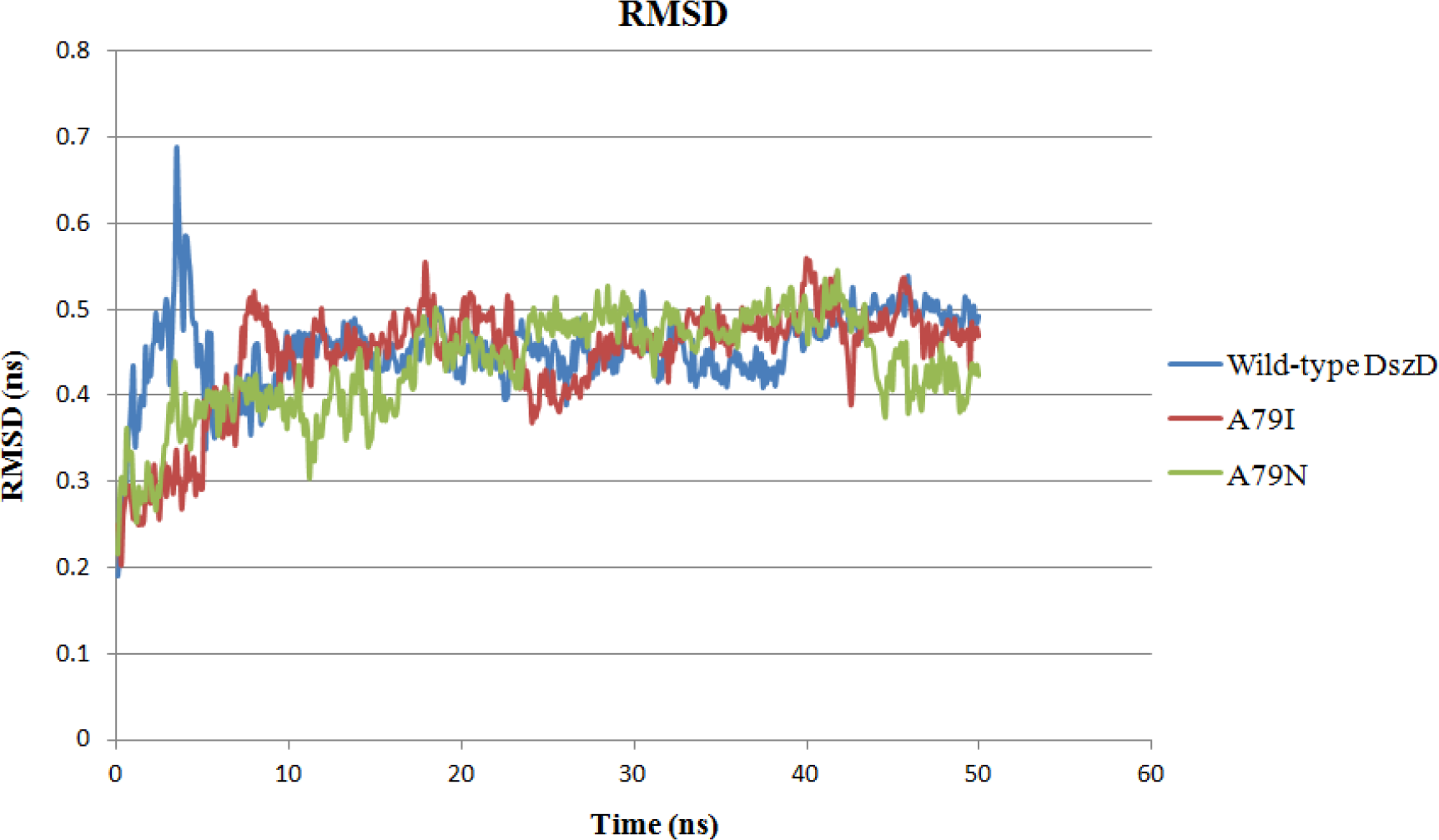
The time evolutions of root-mean-square deviations (RMSD) of the C-α atoms as a function of time for the wild-type DszD and its mutants in complex with FMN-NADH substrates

**Fig. 6.**
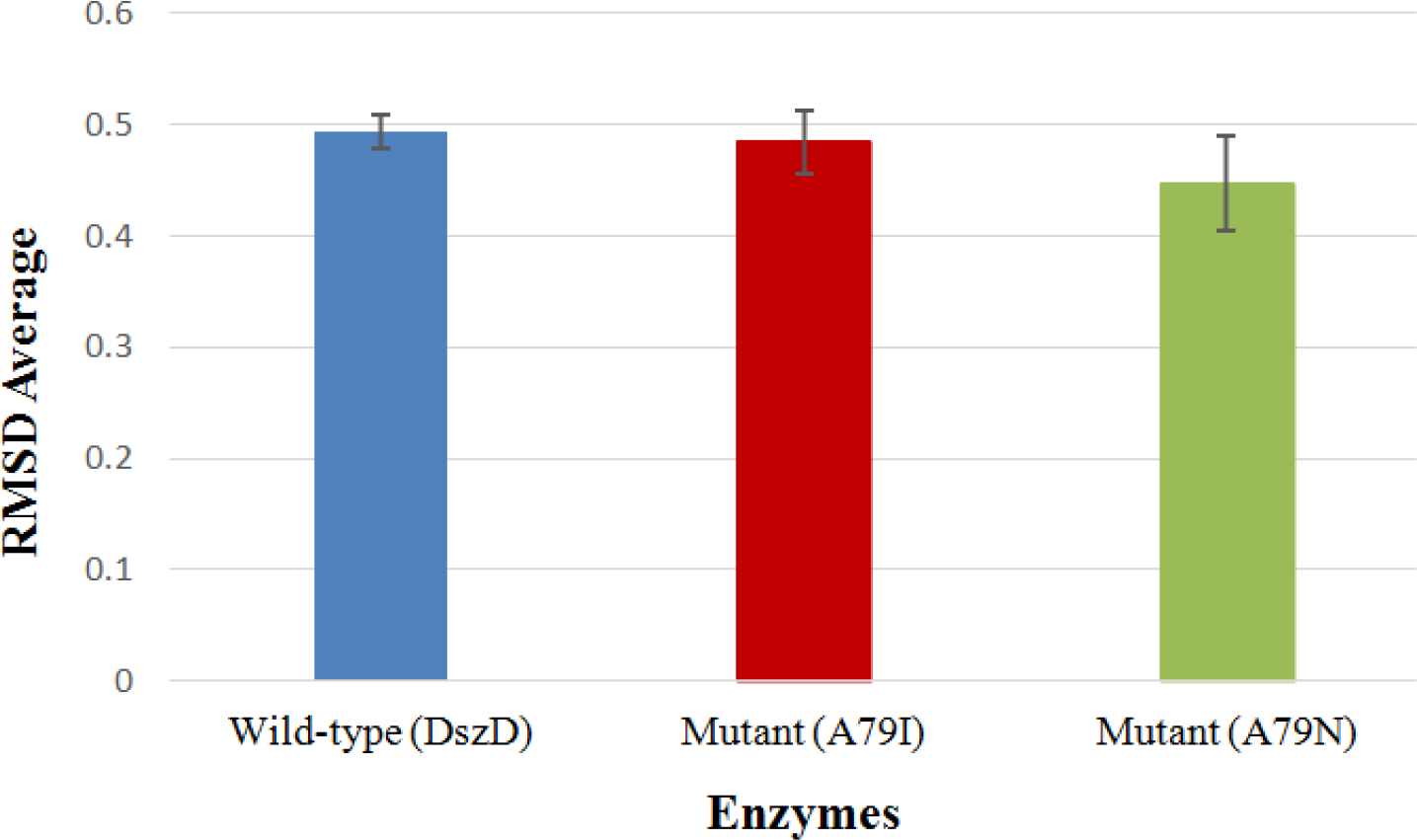
Average of the RMSD of the wild-type DszD, the A79I mutant and A79N in complex with FMN-NADH substrates. The error bar is the standard deviation.

As depicted in Figure 7 the mutation states of the enzyme demonstrate the minimum values of RMSD meaning residues in the whole protein in these two states are fluctuating in a more limited range than wild-type states (DszD) of the enzyme, in other words, the residues are in more stable and equilibrated states (the red and green profiles). The significant reduction in RMSF values is located in binding sites residues 30-45, 80-95 and 165-180.

**Fig. 7.**
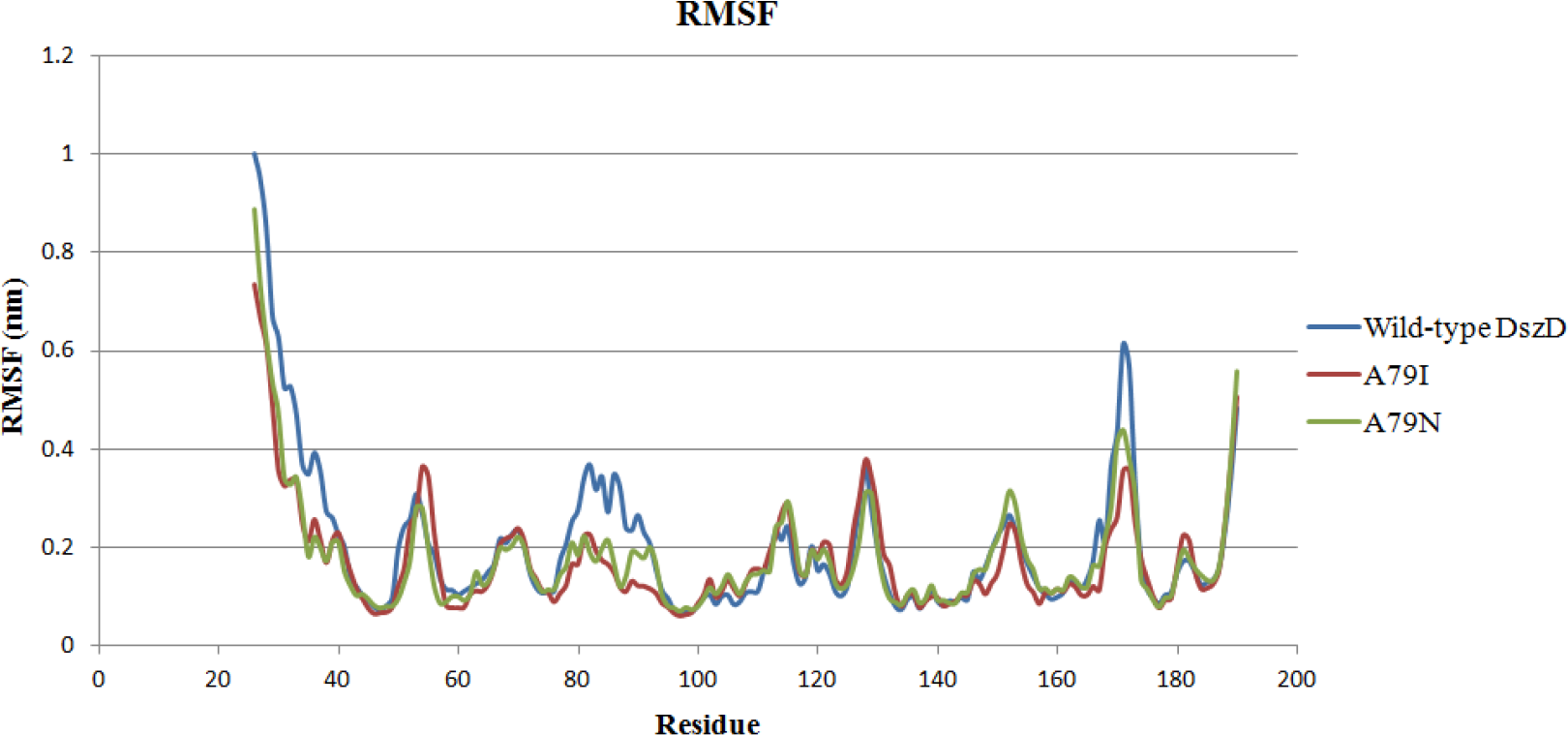
The root-mean-square fluctuation (RMSF) of the C-α atoms as a function of residue for the whole residues of the wild-type enzyme and two mutants.

Dictionary of secondary structure (DSSP) shows the location of residue 79 in the α-Helix region with any change in predicted secondary structure of these mutants. It means that t residue 79 was located in α-Helix region in the wild-type enzyme and mutated enzymes. Consideration of all the available resulting data indicate that replacement of Ala79 with Asn or Ile residue had no effect on predicted secondary structure of these mutants in compare to wild-type enzyme and there was no considerable difference in motifs, regions, active center or steric conformation (Fig. 8). Predicted three dimensional structures of mutants were fully superimposed onto predicted structure of wild-type enzyme and observed their overall folds and maps had highest match and closest similarity to each other.

**Fig. 8.**
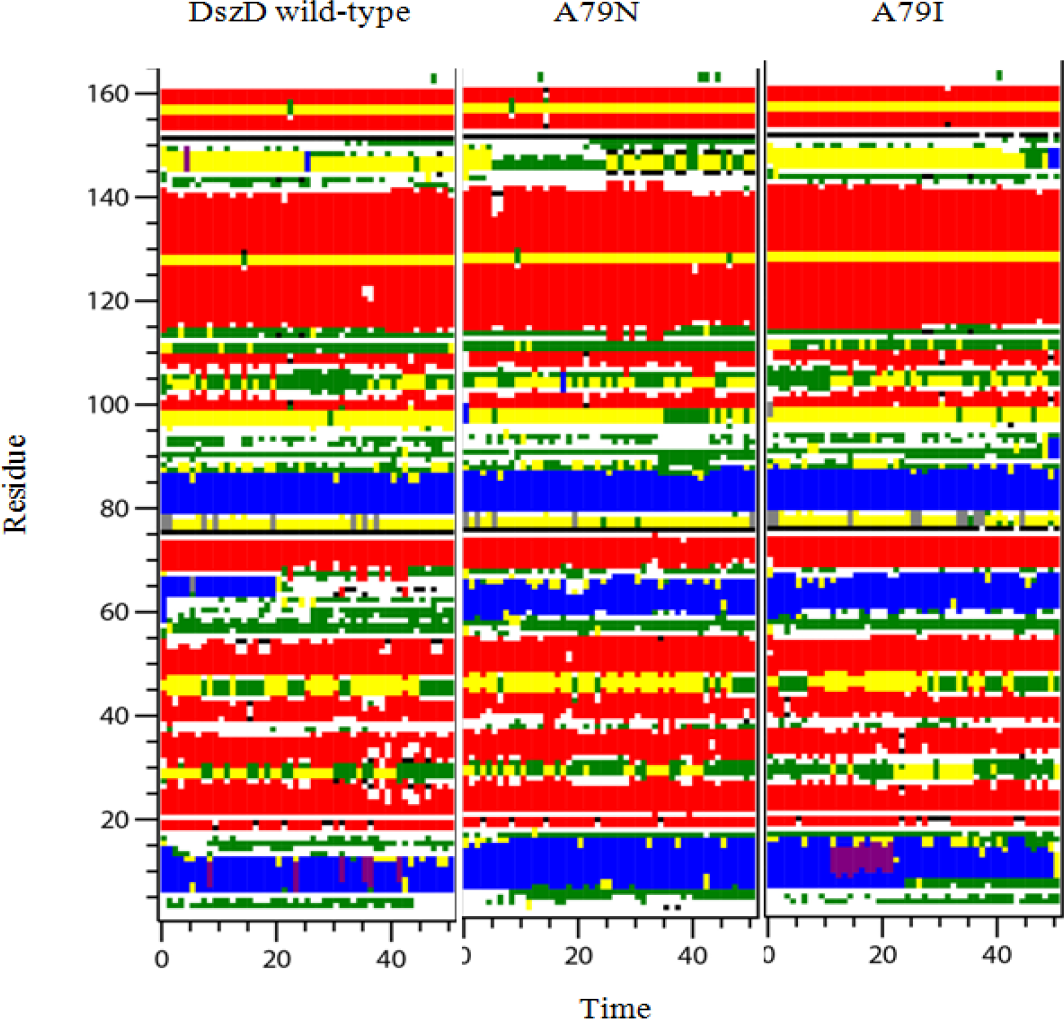
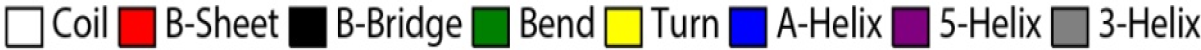
Dictionary of secondary structure (DSSP) of wild-type DszD and Mutants (A79N and A79I).

As indicated in Figure 8 in two mutants of DszD the secondary structures are more preserved than the native state of the enzyme. The helical structure of residues 62 to 67 turned to coils after 20 ns of simulation time in the native state. However, the helical structure of this region is conserved in other mutations to the end of simulation time.

### Hydrogen binding analysis

In comparison with the total number of hydrogen bonds resulted in molecular dockings, the total number of hydrogen bonds in MD simulations has considerably increased due to the equilibrium state of the enzymes in complex with substrates. The biggest number of total hydrogen bonds is obtained for the A79N mutant (Fig. 9).

**Fig. 9.**
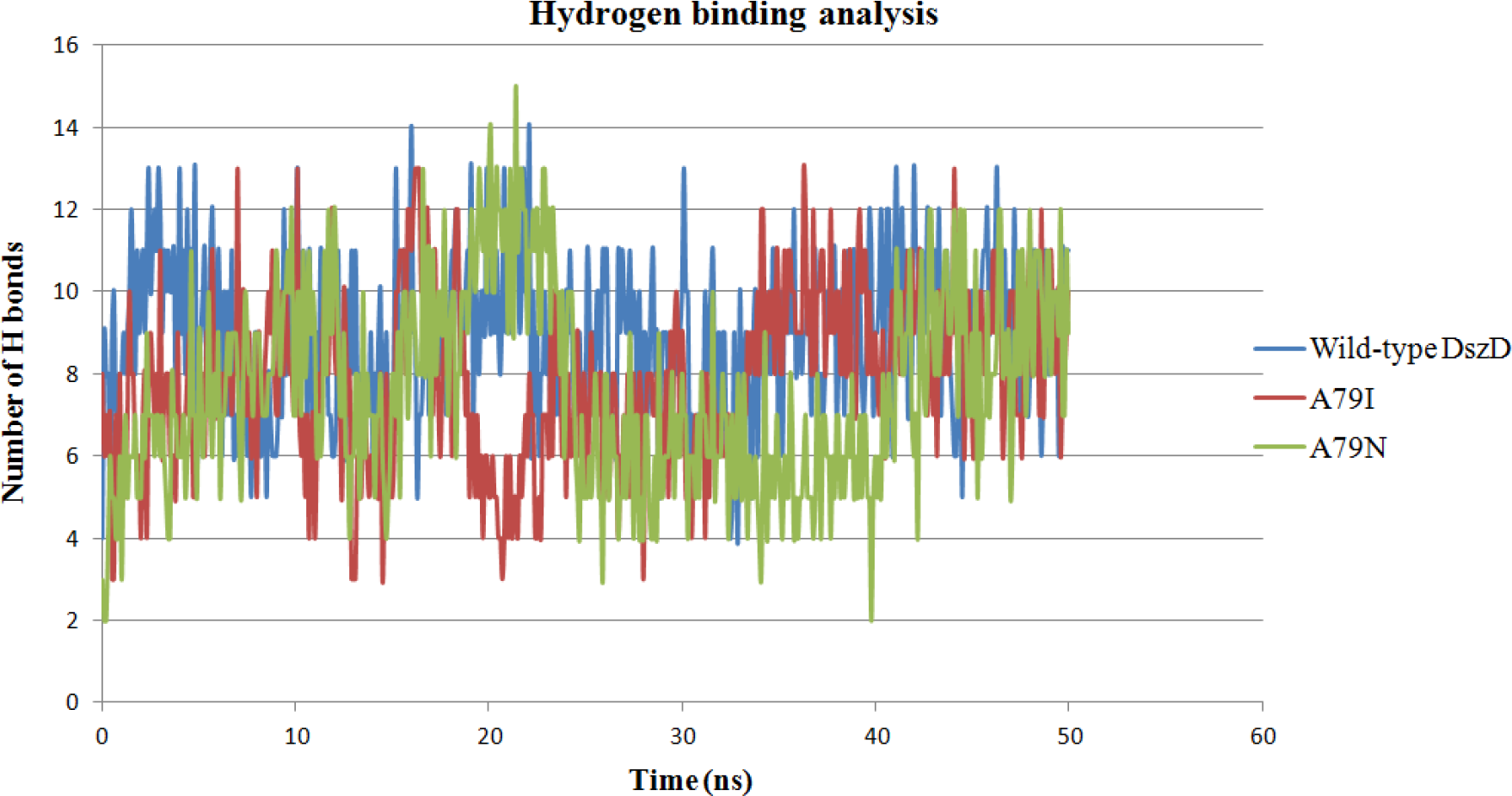
The total number of hydrogen bonds between the enzymes and substrates in all simulation systems.

### Experimental procedures (benchtop operation)

The wild-type *dszD* gene was amplified using the specific primers DszD-F and DszD-R by PCR procedure (Fig. 10A). The *dszD* mutant genes were obtained using the SOEing-PCR by mutagenic primers. The amplicon fragments from the first and second PCRs were purified and used as templates for the third PCR (Fig. 10B, C). The specific primers used were shown in Table 3. The full-length amplicon achieved from the third PCR that was cloned into pET-23a (+) to create recombinant plasmids pET-A79N and pET-A79I. These plasmids carried the DszD mutants A79N and A79I, respectively. The DNA sequencing confirmed presence of the amino acid substitution for each mutant. These plasmids were transfered to *E. coli* BL21(DE3). Then, the transfected bacterial colonies were selected on LB agar plates supplemented with ampicillin.

**Fig. 10.**
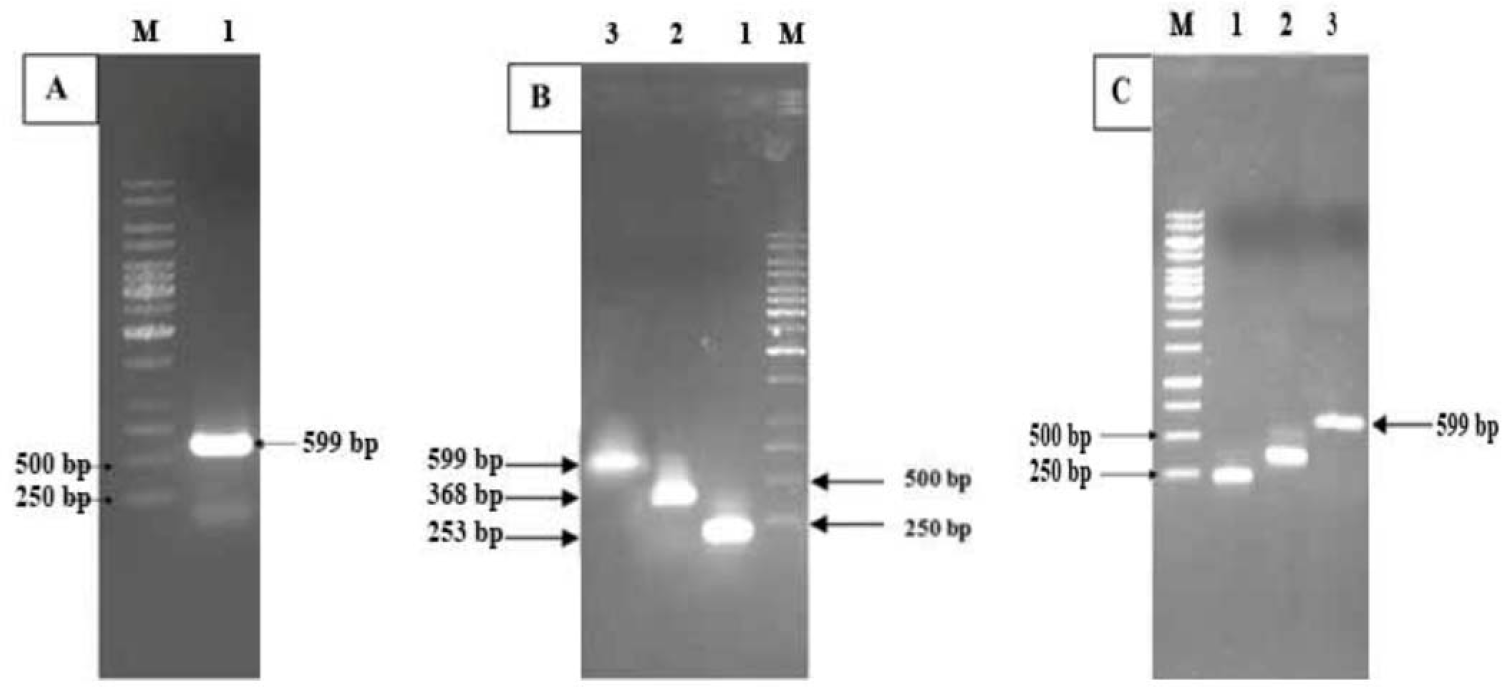
**A** Agarose gel electrophoresis of PCR pattern. Lanes 1 and Lane M show the wild-type *dszD* gene and the DNA Ladder Molecular Weight Marker 1 kb respectively. **B** The SOEing PCR results of the A79I mutant. The line 1, 2 and 3 show the first, second and third PCR amplicon fragments of the A79I mutant respectively. **C** The SOEing PCR results of the A79N mutant. The line 1, 2 and 3 show the first, second and third PCR amplicon fragments of the A79N mutant respectively.

The biochemical analysis of the expression step showed that all of the desired cloned enzymes were overexpressed and had a distinct band on polyacrylamide gel with a calculated molecular mass of 22 kDa (Fig. 11A). Western blot analysis confirmed presence of significant level expression of recombinant wild-type DszD, A79I and A79N mutants (Fig. 11B). The protein concentration was determined by Bradford method (Kruger 2002) with bovine serum albumin as standard. Finally, the enzyme activity of the A79I and A79N mutants were respectively increased 3.4-fold and 5.2-fold compared to the recombinant wild-type enzyme (Table 4). The increased catalytic activity of the A79N mutant was related to the presence of the positively charged carboxamide group of Asparagine (Table 2) that caused to the accelerated electron transfer rate by asparagine 79, which is closer to the isoalloxazine ring than the amine group in alanine 79.

**Table 4.** Comparison of the activity between recombinant wild-type DszD enzyme and its mutants

| Enzyme | Specific activity (U/mg) | compared with the activity of the wild-type enzyme |
| --- | --- | --- |
| Wild-type DszD enzyme | 184 | 1 |
| A79I mutant | 625 | 3.4 |
| A79N mutant | 956 | 5.2 |

**Fig. 11.**
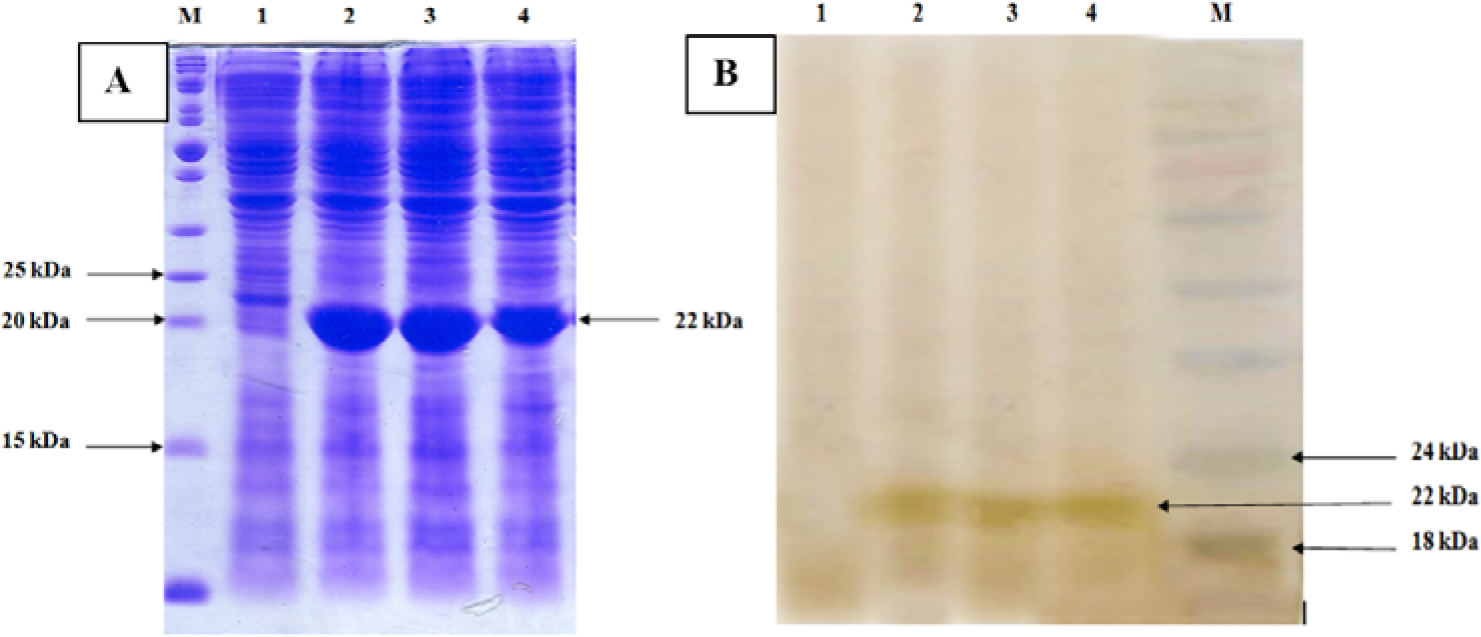
**A** SDS–PAGE analysis of total cell proteins following induction with IPTG of non-transformed *E. coli* BL21, lane 1, and recombinant wild-type *R. erytropolis* IGTS8 DszD, mutant A79N and mutant A79I, Lanes 2, 3 and 4, respectively. Lane M; Protein Molecular Weight Marker. **B** Western-blot analysis using HRP conjugated Anti-His antibody of the recombinants A79N and A79I. Lane 1 is negative control Lanes 2, 3 and 4 are recombinant wild-type DszD, A79N, and A79I after induction with IPTG, respectively. Lane M; Protein Molecular Weight Marker

Also, the increased activity for A79I mutant was higher for recombinant wild-type enzyme.This finding could be attributed to substituting a methyl and ethyl groups for the two hydrogen atoms of the alanine 79. These groups are hydrophobic and have the tendency to fold into active site the enzyme and provide opener active center, allowing the flavin substrate to maneuver much more easily into the proper position. These acquired results reveal a rate-limiting role for alanine 79 in the DszD enzyme. In fact, the presence of the methyl group in alanine side chain in wild-type enzyme has a negative effect in electrons flow of NAD(P)H to flavin by pulling some electrons towards it.

Our results clearly demonstrate that DszD had higher potential to increase the enzyme activity in 4S pathway. Hence, the DszD participation in recombinant homologous or heterologous host cells such as *E. coli* or a recombinant *R. erythropolis* system can cause the better and more efficient biodesulfurization process.

## Conclusion

The bioinformatics and experimental results, revealed the key residues on the active site regions of the DszD enzyme molecule. Among the key residues, the alanine (position 79) had the critical role in the active site of the DszD enzyme. Furthermore, the bioinformatics and experimental results uniformly confirmed the more specific activity of the A79I and A79N mutants compared with the wild-type DszD. Finally, the results of this study can improve the bacterial desulfurization process in the real field in future.

## Acknowledgments

This work was supported by a research grant from the Malek Ashtar University of Technology. We also thank Dr. M.D. Ghafari for advice in this research. Furthermore, the schematic view of the 3-D structure and some analyses were performed with the UCSF Chimera package that was developed by the Resource for Biocomputing, Visualization, and Informatics at the University of California, San Francisco (supported by NIGMS P41-GM103311).

